# In spoken word recognition the future predicts the past

**DOI:** 10.1101/150151

**Authors:** Laura Gwilliams, Tal Linzen, David Poeppel, Alec Marantz

## Abstract

Speech is an inherently noisy and ambiguous signal. In order to fluently derive meaning, a listener must integrate contextual information to guide interpretations of the sensory input. While many studies have demonstrated the influence of *prior* context on speech perception, the neural mechanisms supporting the integration of *subsequent* context remain unknown. Using magnetoencephalography, we analysed responses to spoken words with a varyingly ambiguous onset phoneme, the identity of which is later disambiguated at the lexical uniqueness point^1^. Our findings suggest that primmary auditory cortex is sensitive to phonological ambiguity very early during processing — at just 50 ms after onset. Subphonemic detail is preserved in auditory cortex over long timescales, and re-evoked at subsequent phoneme positions. Commitments to phonological categories occur in parallel, resolving on the shorter time-scale of ~450 ms. These findings provide evidence that future input determines the perception of earlier speech sounds by maintaining sensory features until they can be integrated with top down lexical information.

**Significance statement:** The perception of a speech sound is determined by its surrounding context, in the form of words, sentences, and other speech sounds. Often, such contextual information becomes available *later* than the sensory input. The present study is the first to unveil how the brain uses this subsequent information to aid speech comprehension. Concretely, we find that the auditory system supports prolonged access to the transient acoustic signal, while concurrently making guesses about the identity of the words being said. Such a processing strategy allows the content of the message to be accessed quickly, while also permitting re-analysis of the acoustic signal to minimise parsing mistakes.

---

Typically, sensory input is consistent with more than one perceptual inference, and surrounding context is required to disambiguate. When this ambiguity occurs in a signal that unfolds over time, the system is presented with a critical trade-off: Either prioritise *accuracy* by accumulating sensory evidence over time, or prioritise *speed* by forming interpretations based on partial information. This trade-off is particularly prevalent in speech, which is rife with noise and ambiguity. Further, because language is hierarchically structured, inference occurs both within and across levels of linguistic description: Comprehension of phonemes (e.g. /p/, /b/) is required to understand words; understanding words aids comprehension of their constituent phonemes. How does the human brain strike a balance between speed and accuracy, across these different levels of representation?

When the input is an unambiguous phoneme, low-level spectrotemporal properties are first processed in primary auditory cortex ~50 ms after onset (A1 / Heschl’s gyrus (HG)). Then, higher-level phonetic features are processed in superior temporal gyrus (STG) ~100 ms (Simos et al., 1998; Ackermann et al., 1999; Obleser et al., 2003; Papanicolaou et al., 2003; Obleser et al., 2004; Mesgarani et al., 2014; Di Liberto et al., 2015). These are thought to be purely bottom-up computations performed on the acoustic signal. In natural language, where the acoustic signal is often consistent with more than one phoneme, the system will need to decide which categorisation is the correct one. It is currently unknown where the recognition and resolution of phoneme ambiguity fits relative to this sequence of bottom-up operations.

In order to cope with phoneme ambiguity in speech, the brain uses neighbouring information to disambiguate towards the contextually appropriate interpretation. Most prior research has focused on the use of preceding context, both in terms of the underlying computations and its neural implementation. Concretely, this work suggests that previous context sets up probabilistic expectations about upcoming information, and biases acoustic perception to be consistent with the predicted phonemes (Warren, 1970; Cole, 1973; Samuel, 1981). The left STG and HG appear to be involved in this process, and activity in both regions correlates with the extent to which an expectation is violated (Gagnepain et al., 2012; Ettinger et al., 2014; Gwilliams and Marantz, 2015).

Here we focus on much lesser explored **postdictive** processes, which allow *subsequent* context to bias perception. This phenomenon has been demonstrated behaviourally (Ganong, 1980; Connine et al., 1991; McQueen, 1991; Samuel, 1991; Gordon et al., 1993; McMurray et al., 2009; Szostak and Pitt, 2013), and has been explained in terms of commitment delay: The system waits to accumulate lexical evidence before settling on an interpretation of the phoneme, and maintains sub-phonemic information until the commitment is made. Precisely how the brain implements sub-phonemic maintenance and commitment processes is currently unestablished, but previous research has indicated some likely regions involved. Concretely, activity linked to lexical processing in supra marginal gyrus (SMG) affects phonetic processing in STG at a word’s point of disambiguation (POD) (Gow et al., 2008). The STG and HG have also been implicated in fMRI studies of phoneme ambiguity (Blumstein et al., 2005; Myers and Blumstein, 2008; Kilian-Hutten et al., 2011), in perceptual restoration of masked phonemes (Leonard et al., 2016) and with sensitivity to post-assimilation context (Gow and Segawa, 2009).

In this study we investigate how phoneme perception is influenced by subsequent context, by addressing three questions. First, is the system sensitive to phoneme ambiguity during early perceptual processes, or during higher-order post-perceptual processes? Second, how is sub-phonemic maintenance and phonological commitment neurally instantiated? Third, what temporal constraints are placed on the system — what is the limit on how late subsequent context can be received and still be optimally integrated?

## 2. Materials & Methods

In order to address these questions, we recorded whole-head magnetoencephalography (MEG) across two experiments. In the first experiment, participants listened to syllables that varied along an 11-step continuum from one phoneme category to another (e.g. /pa/ <-> /ba/). Participants classified the sounds as one of the two phoneme categories (e.g. P or B). The syllables provide sensory information about onset phoneme identity but no subsequent context. This protocol is described in detail below.

In the second experiment, a different group of participants listened to items from word <-> non-word continua (” parakeet” <-> “barakeet”). This second set of stimuli thus provides both sensory evidence about the identity of the onset phoneme *as well as* subsequent contextual information. The subsequent information becomes available at the word’s Point of Disambiguation (POD), which refers to the phoneme that uniquely identifies the word being said, and therefore disambiguates the identity of the phoneme at onset. For example, in the word “parakeet” the POD is the final vowel “ee”, because at that point no other English lexeme matches the sequence of phonemes. Therefore, at the POD there is sufficient information in the speech signal to uniquely identify the onset phoneme as /p/. The design of Experiment 2 was inspired by (McMurray et al., 2009).

The first syllables of the words used in Experiment 2 were exactly the same as those used in Experiment 1 — the only difference is that the syllable was followed by silence in the first experiment, and the rest of the word in the second experiment. This allowed us to examine neural responses to the same acoustic signal in isolation and in lexical contexts.

### 2.0 Material creation (common to both experiments)

Word pairs were constructed using the English Lexicon Project (Balota et al., 2007) (ELP), which is a database of phonologically transcribed words and their properties. First, phonological transcriptions of all words beginning with the plosive stops *p, b, t, d, k, g* were extracted. We selected this set of phonemes because it allowed us to examine responses as a function of two phonetic features. Voice onset time (VOT) refers to the amount of time between the release of the stop consonant and the onset of vocal chord vibration. If the amount of time is longer (more than around 40 ms) then the sound will be perceived as voiceless (e.g. *t, p, k*); if the time is shorter (less than around 40 ms) then it will be perceived as voiced (e.g. *d*, *b, g*). Place of articulation (PoA) refers to where in the mouth the tongue, teeth and lips are positioned in order to produce a speech sound. Differences in PoA manifest as spectral differences in the acoustic signal. By measuring responses as a function of both VOT and PoA, we can examine how ambiguity is resolved when it arises from a temporal cue or from a spectral cue, respectively.

Potential word pairs were identified by grouping items that differed by just one phonetic feature in their onset phoneme. For example, the feature voice onset time (VOT) was tested by grouping words with the onset phoneme pairs {t-d, p-b, k-g}, and place of articulation (PoA) was tested with the onset phoneme pairs {p-t, t-k}. Word pairs were selected when they shared 2-7 phonemes after word onset until the phonological sequence diverged. For example, the word pair parakeet/barricade was selected because it differs in voicing of the onset phoneme (p/b), shares the following 4 phonemes (a-r-a-k), and then diverges at the final vowel. This procedure yielded 53 word pairs: 31 differed in VOT and 22 differed in PoA. Words ranged in length from 4-10 phonemes (M=6.8; SD=1.33) and 291-780 ms (M=528; SD=97). Latency of disambiguation ranged from 3-8 phonemes (M=5.1; SD=0.97) and 142-708 ms (M=351; SD=92).

A native English speaker was recorded saying the selected 106 words in isolation. The speaker was male, aged 25, with a Northeast American accent. He said each of the words in a triplet, with consistent intonation (e.g. ↑parakeet,—parakeet, ↓parakeet). The middle token was extracted from the triplet, which promoted similar and consistent intonation and pitch across words. This extraction was done using Praat software (Boersma and Weenink).

Each item pair was exported into TANDEM-STRAIGHT for the morphing procedure (Kawahara et al., 2008; Kawahara and Morise, 2011). In short, the morphing works by taking the following steps: 1) position anchor points to mark the onset of each phoneme of the word pair; 2) place weights on each anchor point to determine the % contribution of each word at each phoneme; 3) specify the number of continuum steps to generate. An explanation and tutorial of the software is available <u>here</u>.

For example, to generate the “barricade” <-> “parricade”, “barakeet” <-> “parakeet” continua shown in Figure 1, anchor points are first placed at the onset of each phoneme in the recorded words “barricade” and “parakeet”, marking the temporal correspondence between the phonemes in the word pair. Next, we decide the amount of morphing to be used at each phoneme in order to generate the unambiguous words/non-words at the end points of the continua. At the first phoneme, the anchor points are weighted as either 100% “barricade” to generate an unambiguous /b/ at onset, or 100% “parakeet” to generate an unambiguous /p/ at onset. All subsequent phonemes until point of disambiguation (“-arak-”) are weighted with equal contributions of each word (50-50). At and after disambiguation, anchors are again weighted at 100%, either towards “parakeet” for the “parakeet-barakeet” continuum, or towards “barricade” for the “parricade-barricade” continuum.

**Fig. 1.**
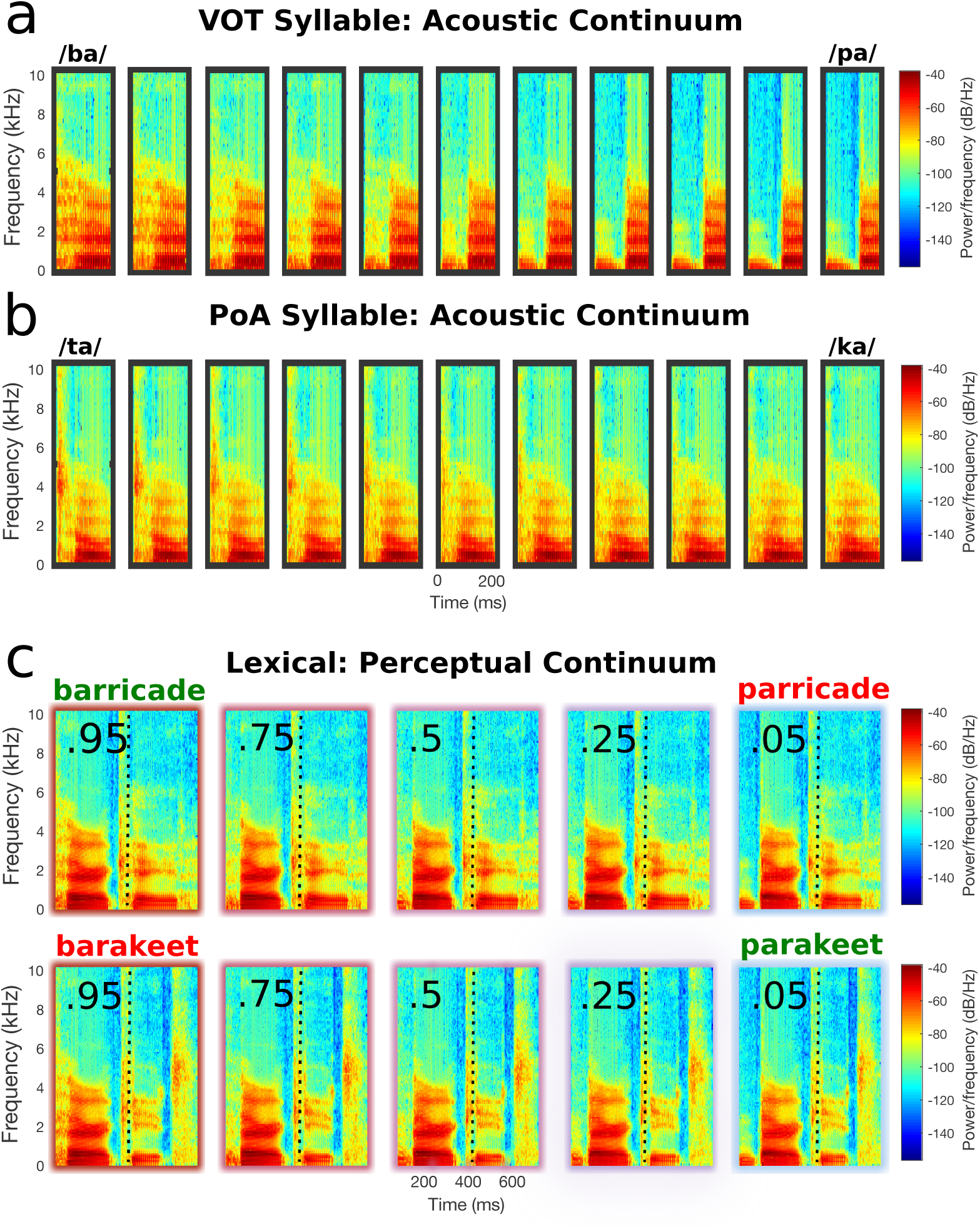
Stimuli examples. **(a)** An example 11-step voice onset time syllable continuum used in Experiment 1. **(b)** An example 11-step place of articulation syllable continuum used in Experiment 1. **(c)** An example 5-step perceptually defined continuum pair used in Experiment 2, generated from the words “barricade” and “parakeet” (shown in green). The resultant non-words “parricade” and “barakeet” are shown in red. The point of disambiguation is represented with a dashed line.

In general, for each pair, all anchor points before the POD are placed at the 50% position, and the first anchor point is positioned either in the congruent position, creating a word (“ parakeet”) or the incongruent (competitor) position, creating a non-word (“ barakeet”). This ensures that apart from the first phoneme, the acoustic signal remains identical across the two word pairs until the disambiguation point. Eleven continua steps were created for each continuum.

The resulting 1166 auditory files were analysed using the Penn Forced Aligner (p2fa) (Rosenfelder et al., 2011) in order to extract the timing of each phoneme’s onset and offset along the length of the word. This created a set of annotation files, which were then visually inspected using Praat (Boersma and Weenink). The accuracy of the p2fa aligner was good overall, but a few manual adjustments were made on approximately 10% of the auditory files in order to ensure correct timing.

### 2.1 Experiment 1

#### 2.1.1 Participants

Twenty-four right handed native English participants took part in the study (11 female; age:*M* =25.44, *SD*=8.44). This sample size was selected based on previous studies using the same MEG machine (e.g. (Gwilliams and Marantz, 2015; Gwilliams et al., 2016)). They were recruited from the New York University Abu Dhabi community and were compensated for their time. All had normal or corrected vision, normal hearing and no history of neurological disorders.

#### 2.1.2 Stimuli

From the word <-> nonword continua described in *Section 2.0*, we extracted just the first syllable (consonant-vowel sequence). This was done for each of the 1166 items. We then amplitude-normed the extracted files to 70 dB.

#### 2.1.3 Procedure

The syllable stimuli were separated into eleven blocks. Each block consisted of two items from each continuum, with the constraint that each item had to be at least three morphed steps away from its paired counterpart. This resulted in a total of 106 trials per block, and 1166 trials total. The assignment of stimulus to block was different for each of the 24 participants, and was balanced by using a latin-square design. Item order was randomised within each block.

Participants heard each syllable in turn, and had to categorise the sound as one of two options that were displayed on the screen. While participants completed the categorisation task, whole-head MEG was being recorded. The screen was approximately 85 cm away from the participant’s face, while they lay in a supine position.

The experimental protocol was as follows. First, a fixation cross was presented for 1000 ms. Then, the two options appeared in upper case, flanking the fixation (e.g. “B + P”). The syllable was played 500 ms later, and the participant needed to indicate which of the two options best matched the syllable they heard by responding with a button box. The options remained on-screen until a response was made. There was no limit placed on how soon participants needed to respond. At each block interval, participants had a self-terminated break. The background was always grey (RGB: 150, 150, 150). All text was in white (RGB: 0, 0, 0), size 70 Courier font. The experiment was run using Presentation^®^ software (Version 18.0, Neurobehavioral Systems, Inc., Berkeley, CA, www.neurobs.com). The recording session lasted ~50 minutes.

### 2.2. Experiment 2

#### 2.2.1 Participants

Twenty-five right handed native English participants took part in the study (15 female; age: *M*=24.84, *SD*=7.3). Six had taken part in Experiment 1 two months earlier. All had normal or corrected vision, normal hearing, no history of neurological disorders, and were recruited from the NYUAD community.

#### 2.2.2 Stimuli

In the second study, we used items from the full word <-> nonword continua. For these items, we wanted to make the onset phonemes across the words differed along a perceptually defined continuum rather than the 11-step acoustically defined continuum used in Experiment 1. In other words, in absence of lexical context, we wanted to make sure the phoneme would be picked out of the pair 0.05, 0.25, 0.5, 0.75 and 0.95 proportion of the time. To set up the materials in this way, we averaged the psychometric functions over subjects for each 11-step continuum used in Experiment 1, and selected the 5 steps on the continuum that were closest to the desired selection proportions (see Figure 2). This converted the continuum from being defined along 11 acoustically-defined steps to being defined along 5 perceptually-defined steps. Continua were removed if the unambiguous endpoints of the continuum were not categorised with at least 80% accuracy for all subjects, or if the position of the ambiguous token was not at least three points away from either end-point of the continuum. This resulted in 49 remaining word pairs, and 490 trials total. These words were amplitude-normed to 70 dB.

**Fig. 2.**
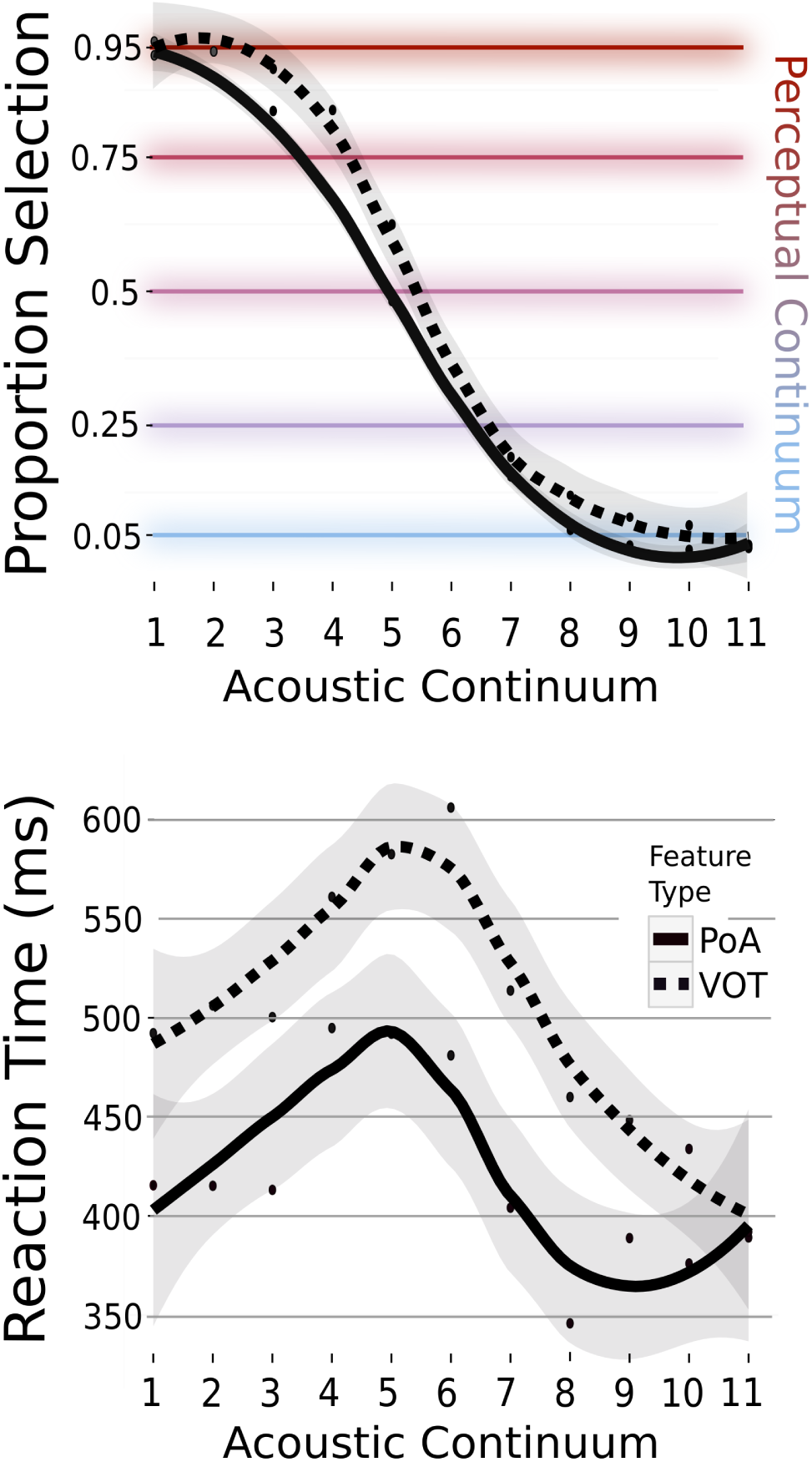
Behavioural results for Experiment 1. Above: Behavioural psychometric function of phoneme selection as a function of the 11-step acoustic continuum. Place of articulation (PoA) and voice onset time (VOT) continua are plotted separately. The coloured horizontal lines correspond to the five behavioural classification positions used to define the perceptual continuum used in Experiment 2. Below: Reaction times as a function of the 11-step continuum; note the slow-down for ambiguous tokens and slower responses to items on the VOT continuum as compared to the PoA continuum.

#### 2.2.3 Procedure

Participants performed an auditory-to-visual word matching task on 2/5 of the auditory items. They were not required to explicitly make judgements about the identity of the onset phoneme. The visual word was either the same as the auditory word (e.g. parakeet-parakeet would require a “match” response) or it was the other word of the pair (e.g. parakeet-barricade would require a “mis-match” response). One item of each five-step continuum was made into a “match” trial (1/5) and one other was a “mis-match” trial (1/5). These conditions were pseudo-randomly assigned using a latin-square procedure. The experiment was split into 5 blocks, and only one token from each continuum appeared in each block. The assignment of item to block was also pseudo-randomised in a latin-square fashion. This resulted in 25 unique experimental orders, across which, items were matched for block order, and match-mismatch assignment.

The experimental protocol was as follows. First, a fixation cross was displayed for 500 ms. Then, while the fixation was still on the screen, the auditory word was presented. If it was a task trial, the visual word appeared 500 ms after the auditory word offset, and remained on screen until participants made a match (left button) or mis-match (right button) decision with their left hand. If it was a no-task trial (3/5 of trials), a blank screen was presented and participants could move to the next trial by pressing either button. The recording lasted ~40 minutes. The apparatus and experiment presentation software was the same as we used in Experiment 1.

### 2.3 Data processing (common to both experiments)

All participants’ head shapes were digitised using a hand-held FastSCAN laser scanner (Polhemus, VT, USA) to allow for co-registration during data preprocessing. Five points on each participant’s head were also digitised: just anterior of the left and right auditory canal, and three points on the forehead. Marker coils were later placed at the same five positions to localise each participant’s skull relative to the sensors. These marker measurements were recorded just before and after the experiment in order to track the degree of movement during the recording.

Stimuli were presented binaurally to participants though tube earphones (Aero Technologies).

MEG data were recorded continuously using a 208 channel axial gradiometer system (Kanazawa Institute of Technology, Kanazawa, Japan), with a sampling rate of 1000 Hz and applying an online low-pass filter of 200 Hz.

MEG data from the two experiments underwent the same pre-processing steps. First, the continuous recording was noise reduced using Continuously Adjusted Least Squares Method (CALM: (Adachi et al., 2001)), with MEG160 software (Yokohawa Electric Corporation and Eagle Technology Corporation, Tokyo, Japan). The noise-reduced data, digital scan and fiducials, and marker measurements were exported into MNE-Python (Gramfort et al., 2014). Bad channels were removed through visual inspection. Independent Component Analysis (ICA) was computed over the noise-reduced data using FastICA in MNE-Python. Components were removed from the raw recording if they contained ocular or cardiac artefacts, which were identified based on the topography of magnetic activity and time-course response. The data were then epoched from 500 ms pre-syllable onset to 1000 ms post-syllable onset for Experiment 1, and 500 ms pre-phoneme onset to 1000 ms post-phoneme onset, for every phoneme in Experiment 2. How we determined the timing of each phoneme is described in section 2.0, last paragraph. Any trials whose amplitude exceeded a +/- 2000 femto-tesla absolute or peak-to-peak threshold were removed. Baseline correction was applied to the epoch using the 200 ms preceding syllable/word onset.

In order to perform source localisation, the location of the subject’s head was co-registered with respect to the sensory array in the MEG helmet. For subjects with anatomical MRI scans (n=4), this involved rotating and translating the digital scan to minimise the distance between the fiducial points of the MRI and the head scan. For participants without anatomical scans, the FreeSurfer “fsaverage” brain was used, which involved first rotation and translation, and then scaling the average brain to match the size of the head scan.

Next, a source space was created, consisting of 2562 potential electrical sources per hemisphere. At each source, activity was computed for the forward solution with the Boundary Element Model (BEM) method, which provides an estimate of each MEG sensor’s magnetic field in response to a current dipole at that source. The inverse solution was computed from the forward solution and the grand average activity across all trials. Data were converted into noise-normalised Dynamic Statistical Parameter Map (dSPM) units (see (Dale et al., 2000)), employing an SNR value of 2. The inverse solution was applied to each trial at every source, for each millisecond defined in the epoch, employing a fixed orientation of the dipole current that estimates the source normal to the cortical surface and retains dipole orientation.

## 3 Results

All results reported here are based on mass univariate analyses. We focused on the following four variables: i) *Acoustics* refers to the item’s position along the 11-step acoustic continuum for Experiment 1, and the 5-step perceptual continuum for Experiment 2. ii) *Ambiguity* refers to how close the item is (measured in continuum steps) to the perceptual boundary between phonological categories. Here we define the perceptual boundary as the position on the continuum where, on average, participants were equally likely to classify the phoneme as one category or the other. iii) *VOT* refers to whether the phoneme was behaviourally classified as voiced or voiceless. iv) *PoA* refers to whether the phoneme was behaviourally classified as being articulated as a bilabial, labiodental or velar stop. We also included *Feature Type*, which refers to whether the phonetic feature being manipulated along the continuum is place of articulation (PoA) or voice onset time VOT).

### 3.1 Behavioural

To analyse behavioural responses in Experiment 1, we applied a mixed effects regression analysis using the *lme4* package (Bates et al., 2014) in *R* (R Core Team, 2012). We included the above four variables as fixed effects and by-subject slopes, as well as Feature Type, the interaction between Feature Type and Ambiguity, and Feature Type with Acoustics. The same model structure was used to fit the reaction time data and the selection data. To assess the significance of each variable, we removed each variable in turn as a fixed effect (but keeping it as a by-subject slope) and compared the fit of that model to the fit of the full model.

For reaction time, we observed a significant effect of Ambiguity, such that responses were significantly slower for more ambiguous items (*χ^2^* = 141.57, *p* <.001). The effect of Acoustics was not significant (*χ^2^* = 3.32, *p* = .068). There was a significant effect of Feature Type, such that responses were significantly slower for VOT continua than PoA continua (*χ^2^* = 99.98, *p* < .001). Ambiguity and Feature Type revealed a significant interaction (*χ^2^* = 8.93, *p* = .002). There was no interaction between Feature Type and Acoustics.

A logistic regression was applied to behavioural selection with the same model structure and model comparison technique. Acoustics was a significant predictor (*χ^2^* = 623.26, *p* <. 001), as well as Feature Type (*χ^2^* = 21.53, *p* < .001). The effect of Ambiguity was not significant (*χ^2^* = 0.68, *p* = .41). Neither was the interaction between Feature Type and Ambiguity (*χ^2^* =2.5, *p* = .11) or Feature Type and Acoustics (*χ^2^* =2.38, *p* = .12). See Figure 2 for a summary of the behavioural results.

Overall, the behavioural analysis indicates that the stimuli are being perceived as intended — we observe a typical psychometric function and a slow-down in responses for more ambiguous items.

### 3.2 Neural

In order to investigate the underlying neural correlates of retroactive perception, we ran a spatio-temporal permutation cluster analysis over localised source estimates of the MEG data (Holmes et al., 1996; Maris and Oostenveld, 2007). This was applied across Heschl’s gyrus (HG) and the superior temporal gyrus (STG) bilaterally, searching a time-window of 0-200 ms after phoneme onset (corresponding either to syllable onset, word onset or POD onset). We implemented the test by running a multiple regression independently at each specified source and time-point. Spatio-temporal clusters were formed for each variable based on adjacent beta coefficients over space and time. In all analyses we used a cluster forming threshold of *p* < .05, with a minimum of 10 neighbouring spatial samples, and 25 temporal samples. See (Gwilliams et al., 2016) for more details concerning this analysis technique.

In the multiple regression, we simultaneously included the four variables described above: Acoustics, Ambiguity, VOT and PoA. Trials were grouped into phoneme categories based on participants’ average behavioural responses in Experiment 1. Number of trials into the experiment and block number were included in all models as nuisance variables. The same analysis was conducted on both Experiment 1 and 2.

### 3.3.1 Experiment 1: Syllable Onset

In terms of main effects, there was a significant effect of Ambiguity, which formed two significant clusters: one in left Heschl’s gyrus (45-100 ms, *p* < .005) and one in the right STG (105-145 ms, *p* = .029). Acoustics formed a cluster in right Heschl’s gyrus, but it was not significant in the permutation test (40-75 ms, *p* = .125). VOT significantly modulated responses in right STG (85-200 ms, *p* < .001), and PoA in left STG (90-150 ms, *p* < .001). The results for Experiment 1 are displayed in Figure 3A-B.

**Fig. 3.**
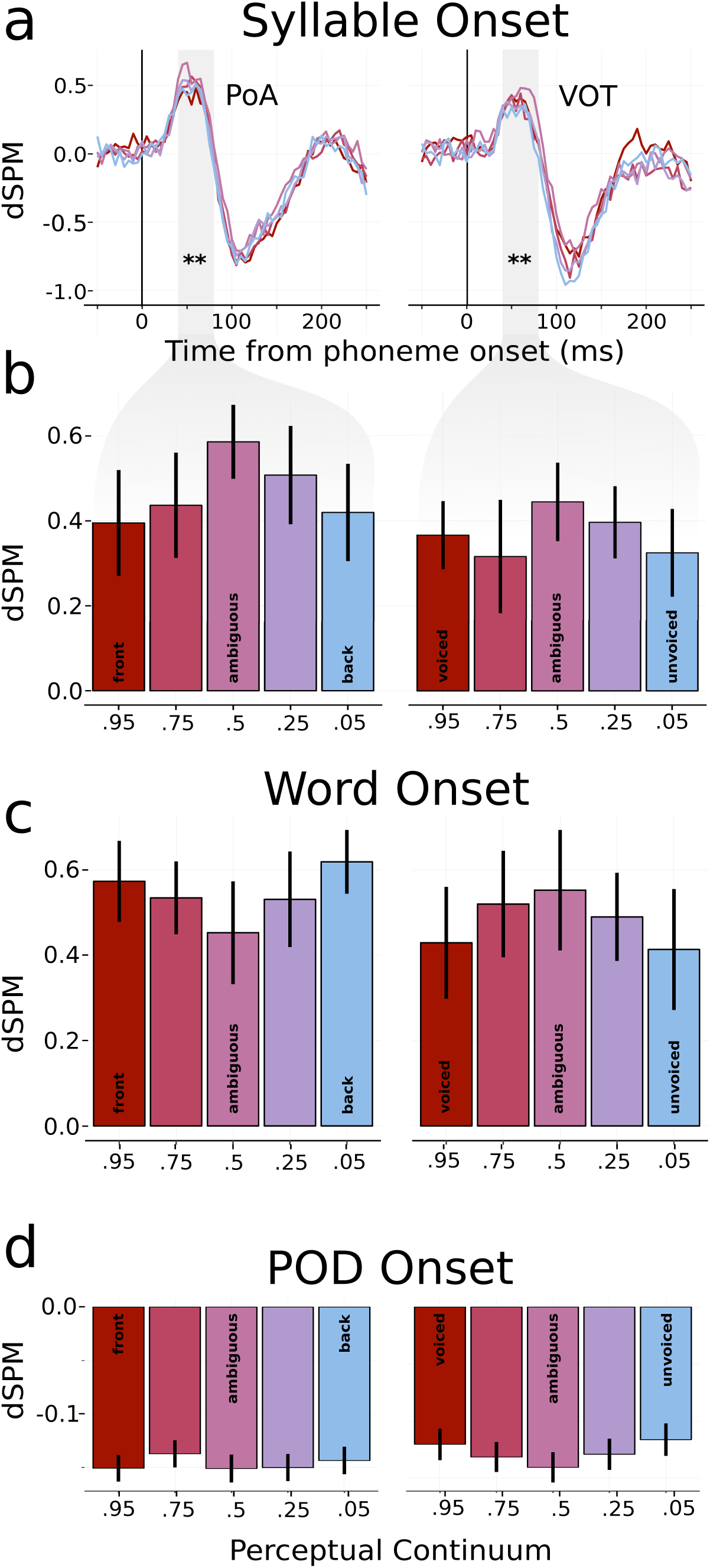
Early responses to ambiguity in left Heschl’s Gyrus (LHG) across the two experiments. **(a)** Timecourse of responses for each ambiguity level averaged over source-localised responses in LHG, plotted separately for place of articulation (PoA) and voice onset time (VOT) continua. **(b)** Averaged responses in LHG over the p50m peak, time-locked to syllable onset in Experiment 1, from 40-80 ms. Note that for the p-t continuum, /p/ is “front” and /t/ is “back”. For the t-k continuum, /t/ is “front” and /k/ is “back”. **(c)** Responses time-locked to word onset, averaged from 40-80 ms. **(d)** Responses time-locked to POD onset, averaged from 40-80 ms. dSPM refers to a noise-normalised estimate of neural activity.

#### 3.3.1 Experiment 1: Acoustic Analysis

Sensitivity to Ambiguity at 50 ms after onset must be reflecting a response to no more than the first 20 ms of the acoustic signal — just the noise burst of the voiceless items, and the initial voicing of the voiced items. To assess what information is available to the system at this latency, we decomposed the first 20 ms of each stimulus into its frequency power spectra using Fast Fourier Transform (FFT). Power at each frequency band from 0-10 KHz, for all stimuli except the fully ambiguous items, was used to train a logistic regression classifier to decode the phonological category (Fig. 4A). Accuracy was significantly above chance level, as determined by 1000 random permutations of phoneme labels (*p* < .001). Accuracy of classification decreased as a function of ambiguity (Fig. 4B), but all continua steps performed greater than chance. Importantly, continua steps themselves could not be decoded from this signal (Fig. 4C), suggesting that this early response indeed scales with distance from the perceptual boundary, and not acoustic properties *per se*. This suggests that the early ambiguity effect we observe in Heschl’s gyrus is not driven by an acoustic artefact generated during the stimuli morphing procedure, for example.

**Fig. 4.**
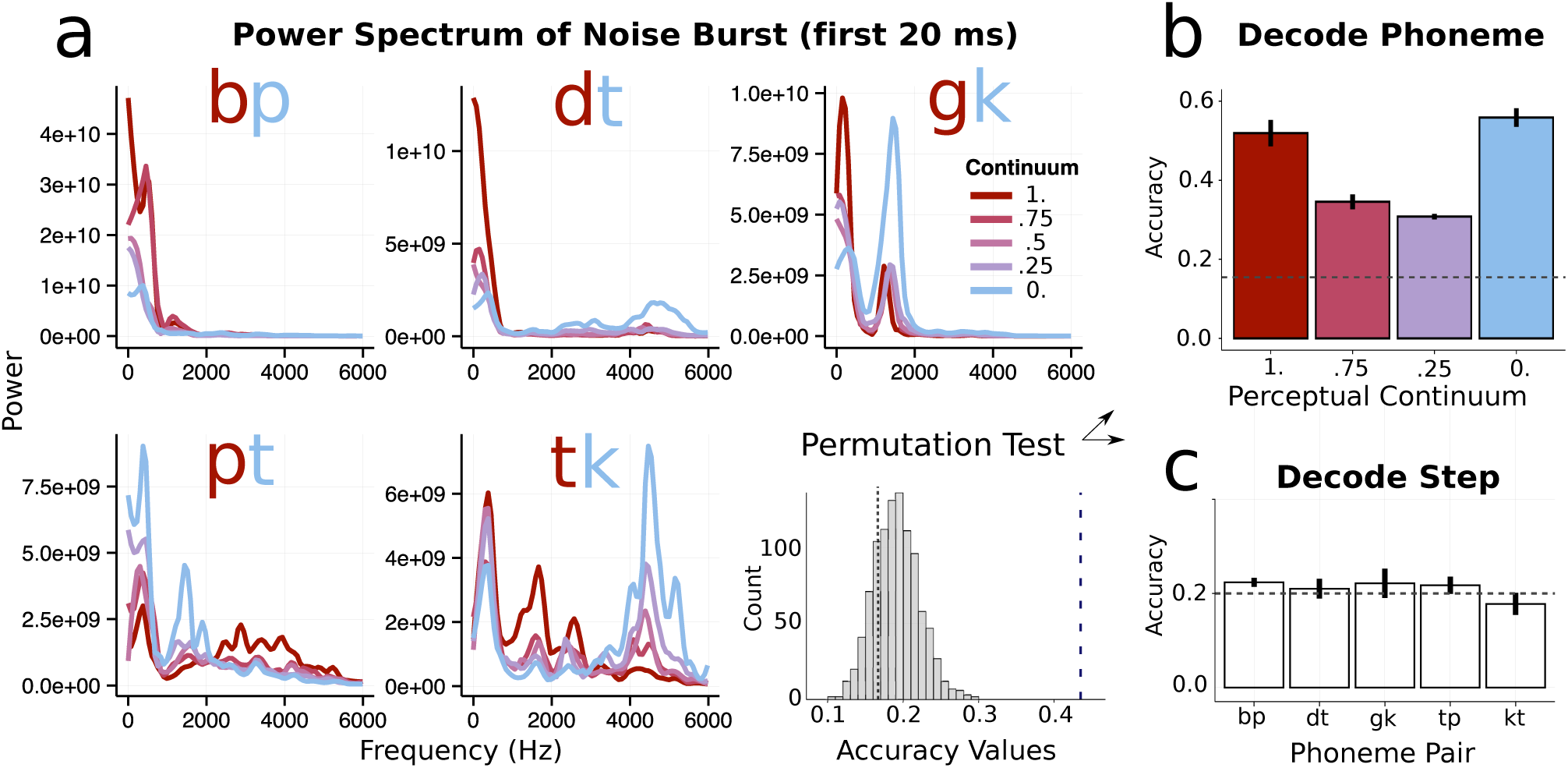
Decoding analysis on acoustic stimuli. **(a):** FFT decomposition of first 20 ms of the auditory stimuli, plotted for each phoneme continuum. The histogram represents the 1000 permutations used to determine the significance of classification accuracy. **(b)**: Accuracy of the logistic regression classifier in identifying the correct phoneme, based on leave-one-out cross validation — accuracy drops off for more ambiguous tokens. **(c)**: Chance-level accuracy in classifying steps along the continuum.

To pursue the stimulus decoding analysis further, we applied the same logistic regression classifier to the first 60 ms of acoustic input – the likely amount of information driving the N100m response (see *Introduction*). The classifier was trained either on a single 60 ms spectral segment of the signal, or three sequential 20 ms spectral chunks. The former provides reasonable spectral resolution but poor temporal resolution; the latter provides the opposite. This novel analysis revealed intuitive results: The classifier more accurately distinguished VOT contrasts (a temporal cue) when trained on three 20 ms chunks, and POA contrasts (a spectral cue) when trained on a single 60 ms chunk. It may be the case that the N100m response is driven by neuronal populations that sample both at fast (~20 ms) and slower (~60 ms) frequencies in order to accurately identity phonemes that vary across each phonetic dimension. This analysis also provides additional support that the early timing of the ambiguity effect is not an artefact, but rather a valid neural response to extant acoustic differences.

#### 3.3.2 Experiment 2: Word Onset

In analysing the results of Experiment 2 we were primarily interested in responses time-locked to two positions in the word. First we will present the results time-locked to word onset, which is also the onset of the phoneme that varies in ambiguity. The analysis was the same as applied for Experiment 1: spatio-temporal cluster test using multiple regression.

In terms of main effects: Ambiguity formed two clusters in left Heschl’s gyrus (150-182 ms, *p* = .034; 144-172 ms, *p* = .063). Acoustics elicited sensitivity in right Heschl’s gyrus (106-152 ms, *p* = .019). Sensitivity to VOT was found in right STG (92-138 ms, *p* < .005); sensitivity to PoA formed two clusters in left STG (86-126 ms, *p* < .005; 88-126 ms, *p* = .028).

The lateralisation of effects observed in Experiment 1 was replicated: Sensitivity to Ambiguity and PoA in the left hemisphere, and to Acoustics and VOT in the right hemisphere. The Ambiguity cluster was identified at ~150 ms in the lexical context, which is later than the effect found for syllable context. However, when looking at the cluster level *t*-values across time (Fig. 5, top left), there was a clear peak in sensitivity to Ambiguity at ~50 ms, too. To test if lexical items also elicit early sensitivity to Ambiguity, we ran a post-hoc mixed-effects regression analysis, averaging just in left Heschl’s gyrus (the locus of the effect in Experiment 1) at 50 ms post word-onset (the peak of the effect in Experiment 1). Ambiguity, Acoustics, Feature Type and their interaction were coded as fixed effects and by-subject slopes. This revealed a significant interaction between Ambiguity and Feature Type (*χ^2^* = 5.9, *p* = .015), and a significant effect of Feature Type (*χ^2^*= 13.14, *p* < .001). When breaking the results down at each level of Feature Type, Ambiguity was a significant factor for PoA contrasts (*χ^2^* = 4.84, *p* = .027) and was approaching significance for VOT contrasts (*χ^2^* = 3.09, *p* = .078). This analysis confirms that the early ambiguity effect is replicated in lexical contexts, albeit with weaker responses. Interestingly, the direction of the effect was reversed for PoA contrasts, whereby more ambiguous tokens elicited less rather than more activity (Figure 3C). This interaction may be due to differences in the task or due to processing syllables versus words. More research would need to be conducted to piece these apart.

**Fig. 5.**
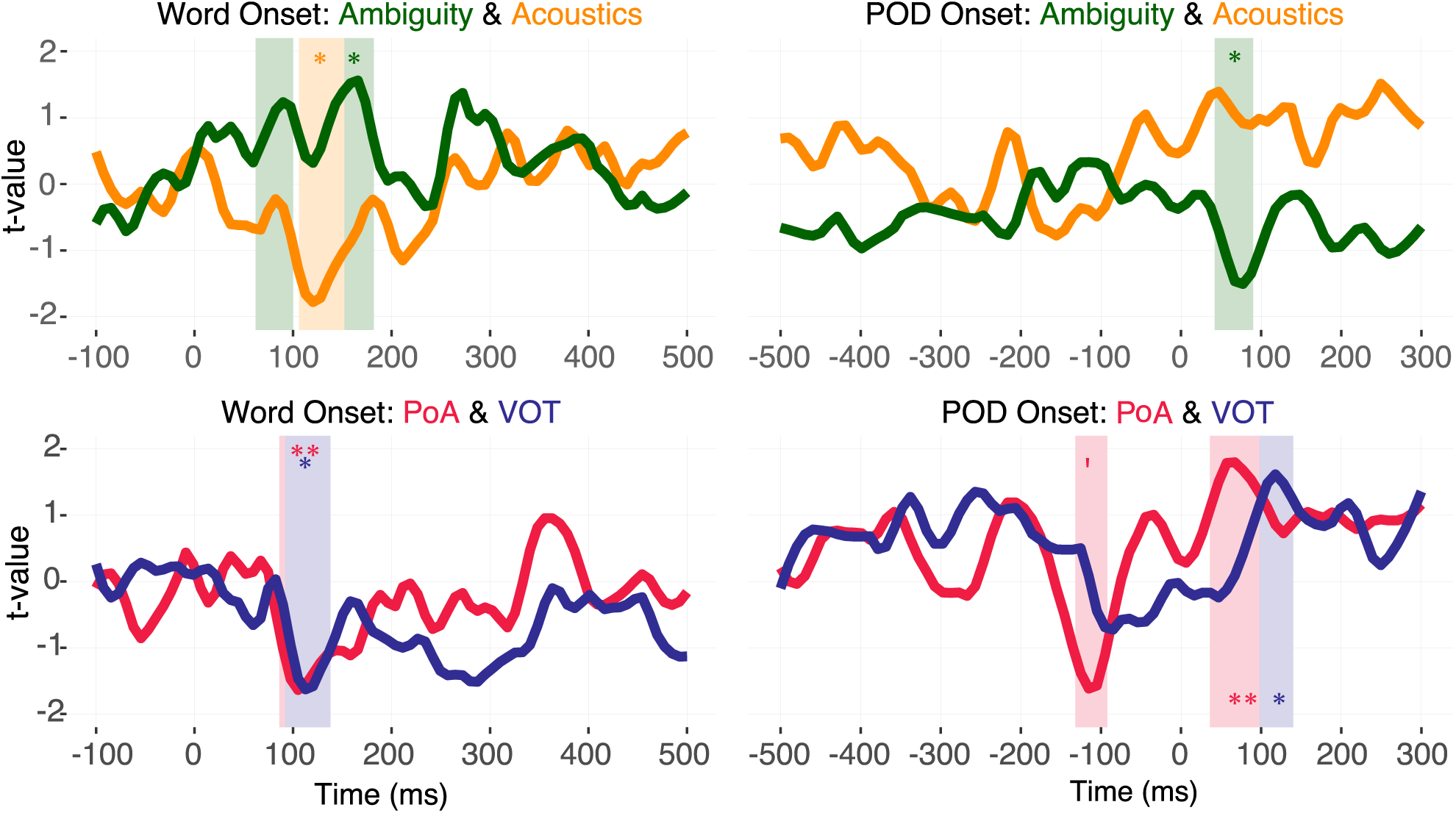
Timecourse of regression analysis for the four primary variables of interest for Experiment 2, time-locked to word onset (left) and point of disambiguation (right). Each trace corresponds to the mean *t*-values averaged in the most significant cluster formed for each variable over time. Note that because the cluster is formed based on the sum of adjacent *t*-values which may be either above 1.96 or below -1.96, the mean value over sources is not directly interpretable as “ *t >* 1.96 = *p*< .05”. Above: *t*-values of Ambiguity and Acoustics variables when put into the same regression model. Below: *t*-values of place of articulation (PoA) and voice onset time (VOT).

#### 3.3.3 Experiment 2: POD Onset

Next we ran the same analysis time-locked to the onset of the word’s point of disambiguation (POD) — this is the phoneme that uniquely identifies what word is being said, and therefore also disambiguates the identity of the phoneme at onset. We used the same analysis technique we used to assess responses at word onset.

In terms of main effects, Ambiguity modulated early responses in left Heschl’s gyrus (50-84 ms, *p* = .011); Acoustics modulated later responses in left Heschl’s gyrus (110-136 ms, *p* = .043). Sensitivity to VOT was found in right STG (98-140 ms, *p* < .01); Sensitivity to PoA was found in left STG (26-96 ms, *p* < .001).

In sum, sensitivity to Ambiguity, Acoustics, PoA and VOT of the onset phoneme is also present at point of disambiguation, with similar lateralisation to that observed at onset (see Fig. 5 & 6).

**Fig. 6.**
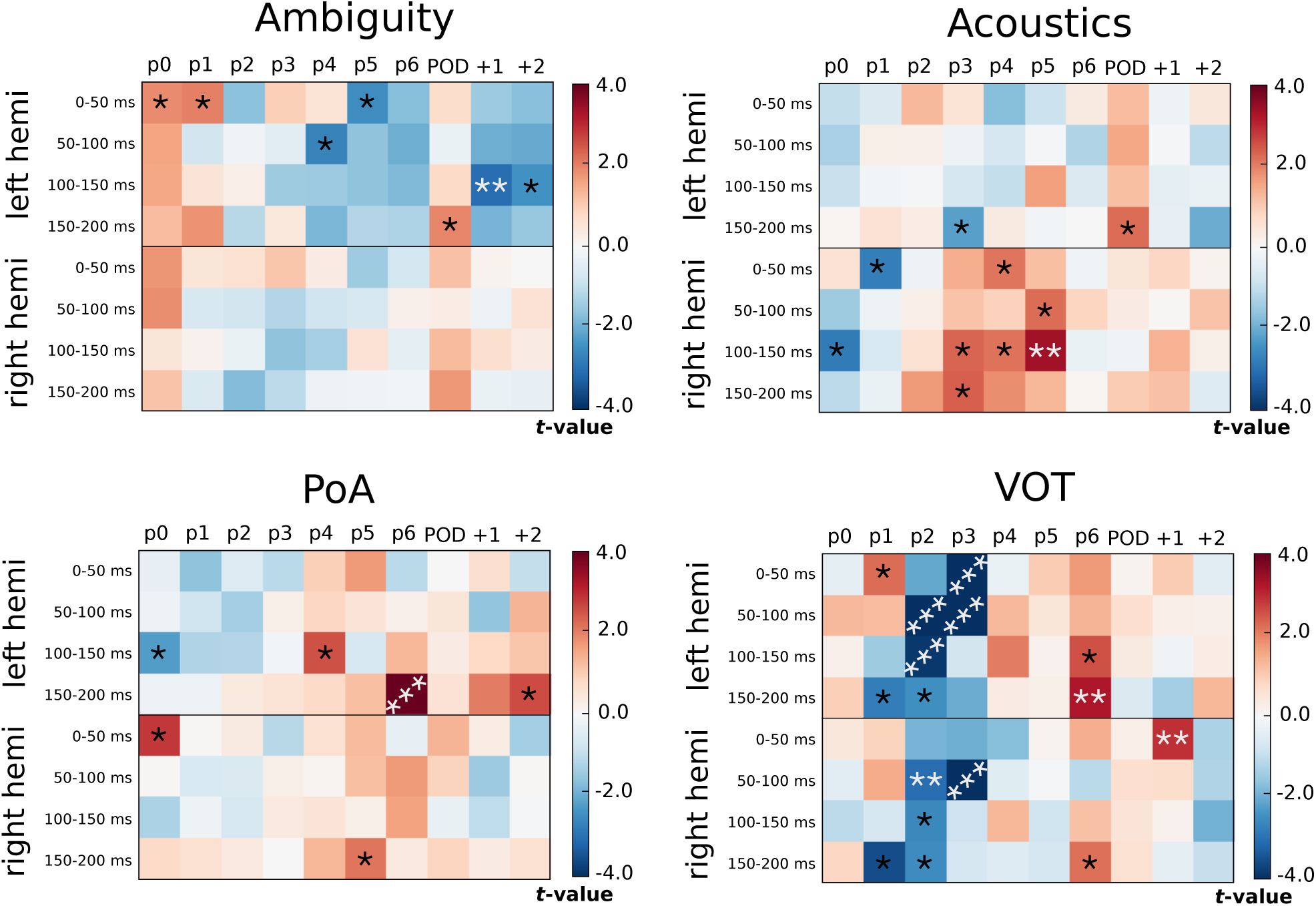
Results of multiple regression applied at each phoneme of the words presented in Experiment 2. Analysis was applied to average source estimates in auditory cortex at different time-windows. For Ambiguity and Acoustics, activity was averaged over left or over right Heschl’s gyrus (the results for both hemispheres are shown). For PoA and VOT, activity was averaged over left or over right superior temporal gyrus and Heschl’s gyrus. The plotted values represent the *t*-value associated with how much the regressor modulates activity in the averaged region and time-window. The analysis was applied separately at the onset of a number of phonemes within the words: p0 = word onset; POD = point of disambiguation; +1 = one phoneme after disambiguation point. Bonferroni corrected *p-*values are shown for reference: * = *p* < .05; ** = *p* < .01; *** = *p* < .001.

#### 3.3.4 Experiment 2: Each Phoneme Onset

Next we wanted to assess whether the re-emergence of sensitivity to the features of the onset phoneme at POD is specific to disambiguation point, or whether it reflected a general re-activation process that could be observed at other positions in the word, too. To test this, we analysed responses time-locked to the 2nd-7th phonemes along the length of the word, as well as the first two phonemes after disambiguation point (Fig. 6).

Spatio-temporal clustering was not the ideal analysis technique to use to test this hypothesis, because statistical strength cannot be assessed if a spatio-temporal cluster is not formed, making it difficult to draw systematic comparisons about the modulation of an effect over time. So instead, we applied the same multiple regression analysis reported above, but simply averaged activity over left or right auditory cortex, and averaged activity within a set of temporal windows. This provided, for each trial, an average measure of neural activity for each hemisphere (2) for each time-window we tested (4) for each phoneme position (10). We corrected for multiple comparisons over these 80 tests using Bonferroni correction. Because the analysis applied here is more conservative than the spatio-temporal test, we can expect some differences in the results reported above for word onset and POD.

The regression was fit to source estimates averaged over just Heschl’s gyrus for Ambiguity and Acoustics, and averaged over both STG and Heschl’s gyrus in the analysis of PoA and VOT. This is because this is where sensitivity to these variables was observed in the responses to syllable onset and word onset.

The results of the analysis are presented in Figure 6, showing the *t*-values and corresponding *p-*values for each multiple regression that was applied at each phoneme, time-window and region. The results show that the re-emergence of sensitivity to each of these variables is not just observed at POD, but also at intermediate positions along the length of the word. There is not a clear relationship between the strength of the reactivation and the phoneme position — for example, the effects do not get systematically weaker with distance from word onset. There are also some differences depending on the feature being analysed: VOT has a particularly strong re-activation at the 3rd and 4th phonemes; PoA and VOT seem to be re-activated bilaterally, whereas Ambiguity remains left lateralised and Acoustics remains primarily right lateralised. These are interesting differences that will require further investigation.

### 3.3.2 Experiment 2: Phonological Commitment

To determine whether the system commits to a phonological category when disambiguation occurs “too late”, we tested for an interaction between disambiguation latency and whether the word resolves to the more or less likely word of the pair given acoustics at onset. The rationale is that if the system commits to a /b/, for example, but then the word resolves to a p-onset word, more effort is required to comprehend the lexical item that was thrown away during the commitment process; however, if no commitment has occurred, there should be a minimal difference between word and non-word resolution because both the cohort of p-onset and b-onset words are still active.

First, we applied the spatio-temporal regression to responses 0-300 ms after to point of disambiguation. Because this question involves higher-level lexical processing, the search area was expanded to include middle temporal gyrus (Fig. 7C). The variables included in the model were lexical resolution (word versus non-word) and its interaction with POD latency, where latency was defined in terms of ms for one test, and in terms of phonemes in the other. No interaction was found between phoneme-defined latency and lexical resolution. However, defining latency in terms of ms did reveal a significant interaction in the left hemisphere between 196-266 ms after POD (*p* = .02). Second, in order to identify the optimal split between “early” and “late”, we averaged activity over the spatio-temporal dimensions of the interaction cluster, and ran a linear mixed-effects regression analysis, testing for an interaction with latency, where latency was shifted incrementally by 1 ms from 200-600 ms after word onset. As can be seen in Fig. 7A, the interaction was maximised when setting the boundary between “early” and “late” between 292-447 ms. When running the same analysis for the Ambiguity variable, no interactions were observed with latency — words that had an ambiguous onset elicited a stronger response at POD regardless of how many ms or phonemes elapsed before disambiguation (Fig. 7D-F).

**Fig. 7.**
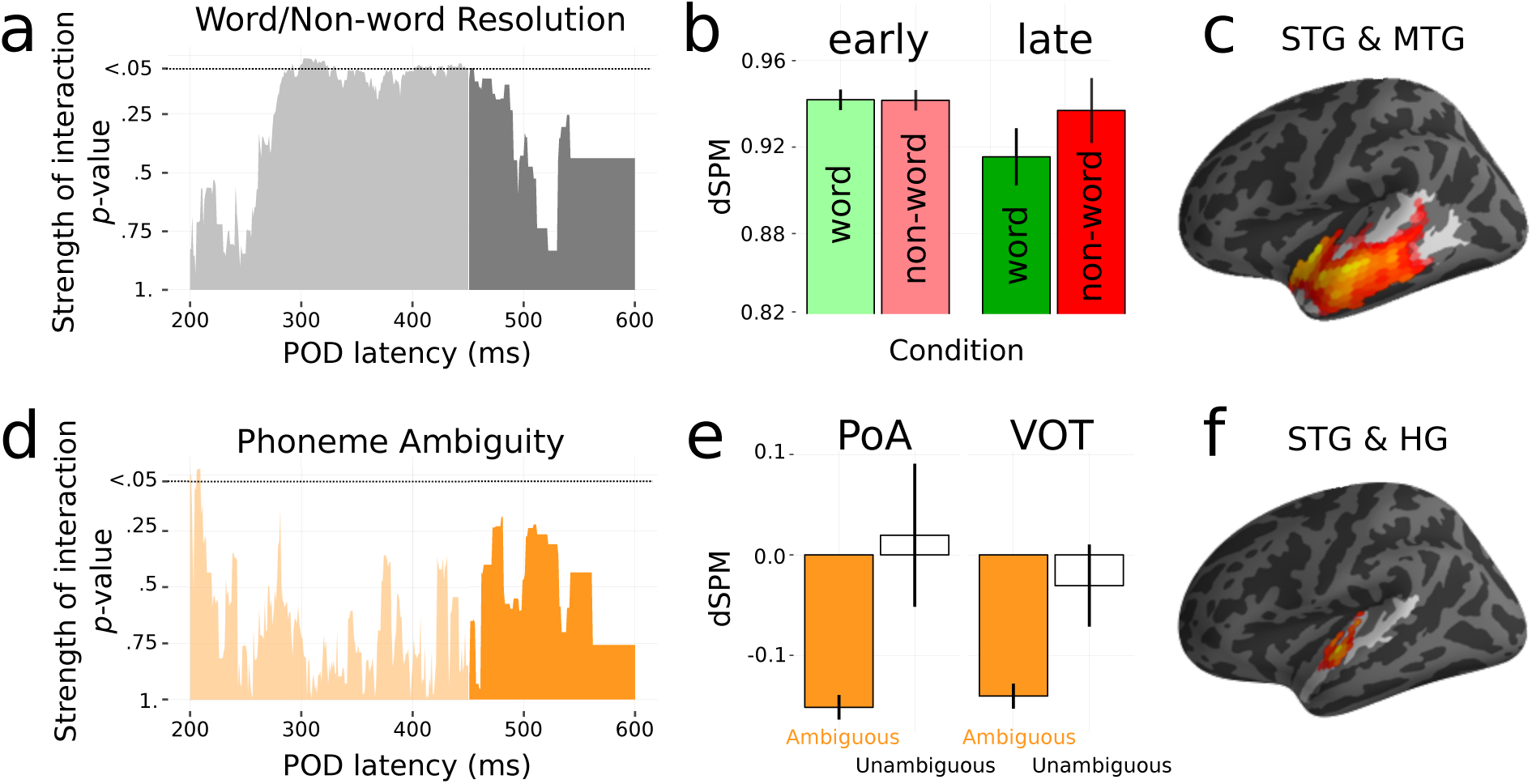
Testing for phonological commitment. Interaction between latency of point of disambiguation (POD) lexical resolution (above) and ambiguity level (below). **(a)** Timecourse of interaction between lexical resolution and “early” versus “late” disambiguation. Early is defined as at or before the increment from word onset shown on the x-axis; late is defined as after the millisecond on the x-axis. The split from light to dark grey shows the final position that the interaction is still significant (450 ms). **(b)** Significant interaction when splitting responses at 450 ms. **(c)** Location of cluster sensitive to the interaction between lexical resolution and latency. **(d)** Non-significant interaction between latency and phoneme ambiguity when splitting latency into “early” vs. “late” incrementally from 200-600 ms post word onset. **(e)** More activity for more ambiguous sounds at disambiguation, even past 450 ms after word onset. **(f)** Location of significant cluster sensitive to ambiguity.

Overall, it appears that non-words are more difficult process than words when disambiguation of the onset phoneme comes later than ms. This suggests that the system does indeed commit to a phonological category after around half a second. The interaction we observe may reflect the system having to re-interpret the input when it has committed to the wrong category (thus perceiving a non-word), or a relative benefit in processing valid words when it has committed to the correct category.

## 4. Discussion

In this study, we aimed to address three research questions. First, does the recognition of phonological ambiguity manifest as an early perceptual process, or a higher-order post-perceptual process? Second, how is sub-phonemic maintenance and phonological commitment neurally instantiated? Third, what temporal constraints are placed on the system — what is the limit on how late subsequent context can be received and still be optimally integrated? We discuss our results in light of these three objectives.

### 4.1 Early sensitivity to ambiguity and acoustics

We found evidence for sensitivity to phonological ambiguity very early during processing, at just 50 ms after onset, in left Heschl’s gyrus. This was orthogonal to sensitivity to position on the continuum, i.e., linear acoustic differences, which was right-lateralised and occurred slightly later. While previous studies have found the p50m to be modulated by VOT (Steinschneider et al., 1999; Hertrich et al., 2000) and PoA (Tavabi et al., 2007), and fMRI studies have found sensitivity to ambiguity in primary auditory cortex (Kilian-Hutten et al., 2011) (see *Introduction*), this is the first evidence of such early responses tracking proximity to perceptual boundaries. This finding supports a hierarchical over reverse-hierarchical processing model (Kilian-Hutten et al., 2011) because sensitivity is apparent before any top-down higher-order influence. This illustrates that early stages of processing are tuned to strikingly complex features of the acoustic signal.

Because of the time it takes the acoustic signal to reach primary auditory cortex, the early ambiguity effect must be reflecting a response to (at most) the first 20 ms of the stimulus. As we were able to decode phoneme category from the spectrotemporal properties of the first 20 ms of the acoustic stimuli (Fig. 4), it is clear that phoneme category information is present in the signal (also see (Blumstein et al., 1977; Stevens and Blumstein, 1978) for a similar conclusion in voiced PoA contrasts). This is consistent with an analysis by synthesis model (Halle and Stevens, 1962; Poeppel and Monahan, 2011) where responses reflect the number of candidate phonemic representations generated by the first ~20 ms of acoustic signal. Neurons fire more when the search space over phonemic hypotheses is large, and less when there are fewer possibilities.

In addressing the first question then, it appears that sensitivity to phonological ambiguity indeed reflects an early perceptual process, and is not driven by top-down influence.

### 4.2 Re-emergence of subphonemic detail

We observed a re-emergence of sensitivity to the acoustics, PoA and VOT of the phoneme heard at onset at each phoneme along the length of the word, at disambiguation point, and at the two phonemes *after* disambiguation. This was specifically time-locked to the onset of each incoming phoneme and was not apparent when analysing based on the time elapsed from word onset (contrast Fig. 5 with Fig. 6). This novel finding is critically important because it supports the hypothesis that the sub-phonemic representation of a speech sound is maintained in superior temporal regions throughout the duration of a word, even while subsequent phonemes are being received; perhaps suggesting that the percept of a speech sound is reassessed at each increment based on the provision of additional input. This finding is also consistent with a recent study using EEG (Khalighinejad et al., 2017), which found evidence for continued maintenance of phoneme-category distinctions.

Further, it appears that phonemic reactivation is a general feature of speech comprehension, rather than a specific mechanism recruited in the presence of ambiguity. Specifically, our results indicate that subphonemic information is maintained even when uncertainty about phoneme identity is low, in two ways. First, re-emergence of phonetic properties was not specific to the ambiguous tokens — it also occurred for the unambiguous phonemes. Second, information about phonetic features continues to be conserved after disambiguating information became available. Overall, these observations are the first to reveal that subphonemic information is maintained, not just in terms of uncertainty about categorisation, but *also* in terms of fine-grained phonetic and acoustic detail of the phoneme under scrutiny. Both sources of information continue to be revisited over long timescales.

In addressing the second question, it appears that commitment delay is instantiated by maintaining phonetic, acoustic and uncertainty information in auditory cortex, and reactivating that information at the onset subsequent phonemes.

### 4.3 Commitment to phonological categories

Finally, we do see evidence for phonological commitment, resolving on a time-scale of ~300-450 ms (see Fig. 7). The superiority of defining latency in terms of elapsed ms rather than phonemes may indicate that commitment is based on the amount of time or number of completed processing cycles rather than intervening information. This process is supported by higher auditory processing regions in anterior STG, a location consistent with a recent meta-analysis of auditory word recognition (DeWitt and Rauschecker, 2012). Critically, this seems to be computed in parallel to the maintenance of subphonemic detail in primary auditory regions. Before ~300 ms there is no cost associated with resolution to a lexical item less consistent with word onset: listeners do not get temporarily mislead (garden-pathed) provided resolution comes early enough (Fig. 7A-B). This suggests that the cohort of words consistent with either phonological interpretation is considered together (e.g., in the presence of b/p ambiguity, both the p-onset and b-onset words are activated). This is fully consistent with previous behavioural studies (Martin and Bunnell, 1981; Gow, 2001; Gow and McMurray, 2007), and a previous eye-tracking study (McMurray et al., 2009), which used similar materials and found look-contingent responses to be dependent upon phonetic information at lexical onset until at least ~300 ms (the longest disambiguation delay they tested). However, after ~450 ms a cost begins to emerge when there is a mismatch between the more likely word given word-onset and the resolving lexical information (e.g., “barricade” is more likely if the onset phoneme was more b-like than p-like, so hearing “parakeet” is a mismatch). This plausibly reflects the recruitment of a repair mechanism, a prediction-error response or re-analysis of the input from making an incorrect commitment.

Finding maintained sensitivity to subphonemic detail in parallel to phonological commitment is very important for the interpretation of psychophysical research, which has implicitly equated insensitivity to within-category variation with phonological commitment (Connine et al., 1991; Szostak and Pitt, 2013; Bicknell et al., 2015). This previous work has largely converged on a processing model whereby phonological commitment can be delayed for over one second after onset. Our results indicate, in contrast, that while subphonemic detail is indeed maintained over large time-scales, this does not implicate that commitment is also put off for this length of time.

In addressing the third question, it seems that subsequent context can be optimally integrated if it is received within around half a second. This is when the system commits to a phonological interpretation. However, subphonemic detail is maintained past the point that the system makes such a commitment.

### 4.4 Relationship to models of speech processing

It is unclear which model of speech processing can account for these data. While Shortlist (Norris, 1994) and Shortlist B (Norris and McQueen, 2008) may be able to model recovery from lexical garden-paths, they do not explicitly model processing of subphonemic detail. While the MERGE model (Norris et al., 2000) is capable of modelling such detail, it proposes no feedback from the lexical to phoneme levels of analysis, which is inconsistent with our observation that (sub-)phonemic representations are reactivated when top-down lexical information becomes available. Although it has been demonstrated that TRACE (McClelland and Elman, 1986) can be modified to simulate recovery by removing phoneme-level inhibition (McMurray et al., 2009), it does not provide the architecture to model initial sensitivity to phoneme ambiguity, or account for how the percept of speech sounds is modulated by past and future linguistic information (see (Grossberg and Kazerounian, 2011) for an overview of TRACE limitations). It is also unclear whether this modification would interfere with TRACE’s success in accounting for a range of observations in spoken word recognition (see (Gaskell, 2007) for a review). One model proposed to deal with TRACE’s shortcoming is Adaptive Resonance Theory (ART): each speech sound produces a resonance wave that is influenced by top-down information until it reaches equilibrium and surfaces to consciousness (Grossberg, 2003). While this theory is consistent with the idea that there is a critical time-limit to receive top-down information, it suggests that there is a linear decay in subphonemic information as temporal distance from the phoneme increases. Our results do not support that conjecture. Instead, they suggest that subphonemic information is re-evoked later in processing, with a similar magnitude as that experienced at onset. In light of the present results, one shortcoming of these models is their attempt to explain spoken word recognition with a single mechanism, built on the assumption that acoustic-phonetic information is lost once a phonological categorisation is derived. Instead, our results suggest that a three-element processing model is more appropriate, allowing for a dynamic interaction between phonetic, phonological and lexical levels of analysis.

### 4.5 Conclusion

Later sounds determine the perception of earlier speech sounds through the simultaneous recruitment of acoustic-phonetic and phonological computational pathways. This facilitates contact with lexical items in order to derive the message of the utterance, as well as continued revisitation to the phonetic level of analysis to reduce parsing errors. In this manner, lexical selection can be achieved rapidly, while also reducing the likelihood of mistakes in phonological segmentation. The human brain therefore solves the issue of processing a transient hierarchically structured signal by recruiting complementary computations in parallel, rather than conceding to the trade-off between speed and accuracy.

## Acknowledgements

This research was supported by ERC-2011-AdG 295810 BOOTPHON, ANR-10-IDEX-0001-02 PSL and ANR-10-LABX-0087 IEC to TL; NIH 2R01DC05660 to DP; NYU Abu Dhabi Institute under grant G1001 to AM. We would like to thank Kyriaki Neophytou for her help with data collection and Lena Warnke for help with stimulus creation.

